# Virtual perturbations to assess explainability of deep-learning based cell fate predictors

**DOI:** 10.1101/2023.07.17.548859

**Authors:** Christopher J. Soelistyo, Guillaume Charras, Alan R. Lowe

**Affiliations:** University College London, London, UK; Institute for the Physics of Living Systems, London, UK; Alan Turing Institute, London, UK

## Abstract

Explainable deep learning holds significant promise in extracting scientific insights from experimental observations. This is especially so in the field of bio-imaging, where the raw data is often voluminous, yet extremely variable and difficult to study. However, one persistent challenge in deep learning assisted scientific discovery is that the workings of artificial neural networks are often difficult to interpret. Here we present a simple technique for investigating the behaviour of trained neural networks: virtual perturbation. By making precise and systematic alterations to input data or internal representations thereof, we are able to discover causal relationships in the outputs of a deep learning model, and by extension, in the underlying phenomenon itself. As an exemplar, we use our recently described deep-learning based cell fate prediction model. We trained the network to predict the fate of less fit cells in an experimental model of mechanical cell competition. By applying virtual perturbation to the trained network, we discover causal relationships between a cell’s environment and eventual fate. We compare these with known properties of the biological system under investigation to demonstrate that the model faithfully captures insights previously established by experimental research.

## 1 Introduction

### 1.1 Deep learning and scientific discovery

Scientific models are typically built to explain events in the natural world. This often consists of building relationships between elements of the world; for example, the relationship between the current environment of a biological cell and its eventual mitotic or apoptotic fate.

The discovery of patterns within observed data is a key strength of deep learning. Therefore, its recent emergence has raised the potential for the automated generation of accurate scientific models [1]. A deep neural network (DNN), through being trained on large amounts of real-world data, can create internal representations that capture relationships in the natural world - in other words, a scientific model that can predict the outcome of experimental observations. This stems from the incredible power of DNNs to approximate input-output mappings given a sufficient corpus of training examples.

Abstractly, the aim of machine learning (ML) in general is to produce a model that can approximate some desired mapping *f* - which represents the natural phenomenon - from an input domain *X* to an output domain *Y* :

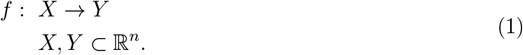

The goal is to learn some function *g* that approximates *f* (i.e., *g*(*x*) *≈ f* (*x*)). This approximate function could be modelled by a DNN.

### 1.2 The problem of scientific explainability

There are at least two key issues with using DL to generate scientific models. The first is that it is notoriously difficult to explain how a deep neural network arrives at its outputs. This is due largely to typical complexity of DNNs, rendering them “black boxes” resistant to human interpretation. Therefore, it can be difficult to build accurate DNNs that are simultaneously explainable. However, we hypothesise that it is possible to extract insights learnt by the DNN by focusing on causal relationships between input and output, a theme that will be further developed in Section 3 within the context of deep-learning based cell fate predictors.

The other key issue with DL-based science is that it may be possible to develop a multitude of models that achieve the same degree of accuracy through a diversity of internal mechanisms. Hence the learnt function, or “theory”, *g* could belong to a wider set *G* of equal-performance functions, what Semenova et al. [2] call a “Rashomon set”. Therefore, by fitting the data, a trained model will not necessarily have learnt underlying natural phenomenon; it will have learnt *a* theory, perhaps one of many, that is consistent with the observed data. We will return to this point in the discussion (Section 4).

### 1.3 DL as a tool in bioimaging

One of the scientific fields most promising for the application of deep learning is cell bioimaging. This is due primarily to the inherent complexity and volume of biological data, as well as its tractability as a problem. The continued development of microscopy methods has made available a wealth of image data related to cell appearance, organisation and behaviour. Meanwhile, automation and cell segmentation technology has further enabled the high-throughput collection of vast amounts of spatio-temporal data, allowing for the capture of time-lapse videos [3].

This abundance of data provides both opportunities and challenges for the cell imaging field. Cell biologists can access an unprecedented amount of information. However, as noted by Ouyang and Zimmer [4], the volume of data produced by modern imaging technologies has “outgrow[n], often vastly, the capacity of manual analyses and human inspection”. As a result, this field has seen a recent explosion of studies applying deep learning [5]. The capability of DNNs to extract patterns from complex data has led to their application by cell biologists in tasks as varied as feature extraction [6, 7, 8, 9], morphology-based classification [10, 11, 12, 13, 14, 15, 16], image segmentation [17, 18, 19, 20, 21, 22, 23, 24], synthetic data generation [25, 26] and more.

In recent years, the field has also seen applications of explainable deep learning to extract scientifically relevant patterns from complex bioimaging data. For example, Zaritsky et al. [27] use an autoencoder combined with linear discriminant analysis to identify morphological features indicative of metastatic efficiency level in melanoma cells. We previously developed a framework to identify biophysical determinants of cell fate in cell competition [28]. The present study aims to extend our previous work, by systematically examining input-output relationships in a DNN model designed to predict cell fate.

## 2 Model system: predicting cell fate using a deep neural network

Our model system is a competition between wild-type MDCK cells (MDCK^WT^) and scribble-knockdown MDCK cells (scrib^kd^), where the latter expresses an shRNA that silences expression of the Scrib gene. The loss of the scribble polarity protein is a deleterious mutation, and triggers competitive interactions between the MDCK^WT^ and scrib^kd^ cells, where the presence of the former leads to elimination of the latter via apoptosis.

This is an example of cell competition, a biological phenomenon that involves the specific elimination of a particular cell population by competitive interactions with one or more other cell populations within a tissue [29]. The ability of a cell to compete in this fashion is called its “fitness” [30]. A hallmark of this phenomenon is that the less-fit (“loser”) cells are viable when cultured alone but are eliminated when they come into contact with the fitter (“winner”) cells. In many systems, cell competition acts a quality-control mechanism [31, 32, 33, 34, 35].

A decade of experimental research has revealed the dynamics of MDCK^WT^ and scrib^kd^ cell competition [36, 37, 38, 39]. MDCK^WT^ cells have higher homeostatic tension than scrib^kd^ cells, therefore, when the populations are put in contact, the presence of MDCK^WT^ cells induces crowding in scrib^kd^ cells, which leads to their mechanical compression and eventual elimination via apoptosis. The key role of crowding in fate determination in scrib^kd^ cells was then independently rediscovered by a deep neural network trained to predict cell fate (apoptosis vs. mitosis) based on time-lapse videos of single cells taken by fluorescence microscopy [28].

In our previous study, we built a three-stage neural network architecture (referred to as a *τ* -VAE), to predict cell fate from a sequence of microscopy images. In particular, the network converts a time-lapse video portraying scrib^kd^ cell throughout its lifetime, called a “trajectory”, into a prediction of the cell’s eventual fate. Each trajectory was *∼*8.5 hours in duration, and each video frame captures a 21.3 *×* 21.3*μ*m area. These trajectories were obtained by tracking the U-Net [17] segmentations of each cell, using the *btrack* package [40, 21]. In each video, MDCK^WT^ and scrib^kd^ nuclei are distinguished by tagging the Histone H2B protein with different fluorophores, green fluorescent protein (GFP) for MDCK^WT^ and red fluorescent protein (RFP) for scrib^kd^. Hence, the *τ* -VAE receives information regarding both cell type and nuclear morphology. Importantly, the videos do not contain any evident morphological changes of the nucleus prior to the fate event (e.g. transition to prometaphase prior to cell division). This ensures that the model is trained to *predict* cell fate, rather than simply *observe* morphological indicators of cell fate.

The first stage of the *τ* -VAE is the encoder part of a variational autoencoder [41, 42, 43], which converts each image into a latent vector representation. The second is a projective transform that maps points in the *β*-VAE latent space to points in its principal component (PC) space, learnt by applying principal component analysis (PCA) to a large set of images. The third is a temporal convolutional network (TCN) [44] that receives the PC representation of the input video, then outputs the cell fate prediction (**Fig. 1**). It was found that the PC-transform step was necessary to obtain embedded representations that were interpretable. The results were most striking for PC0 and PC1, which represented cell type and cell size/nuclear area respectively.

**Figure 1.**
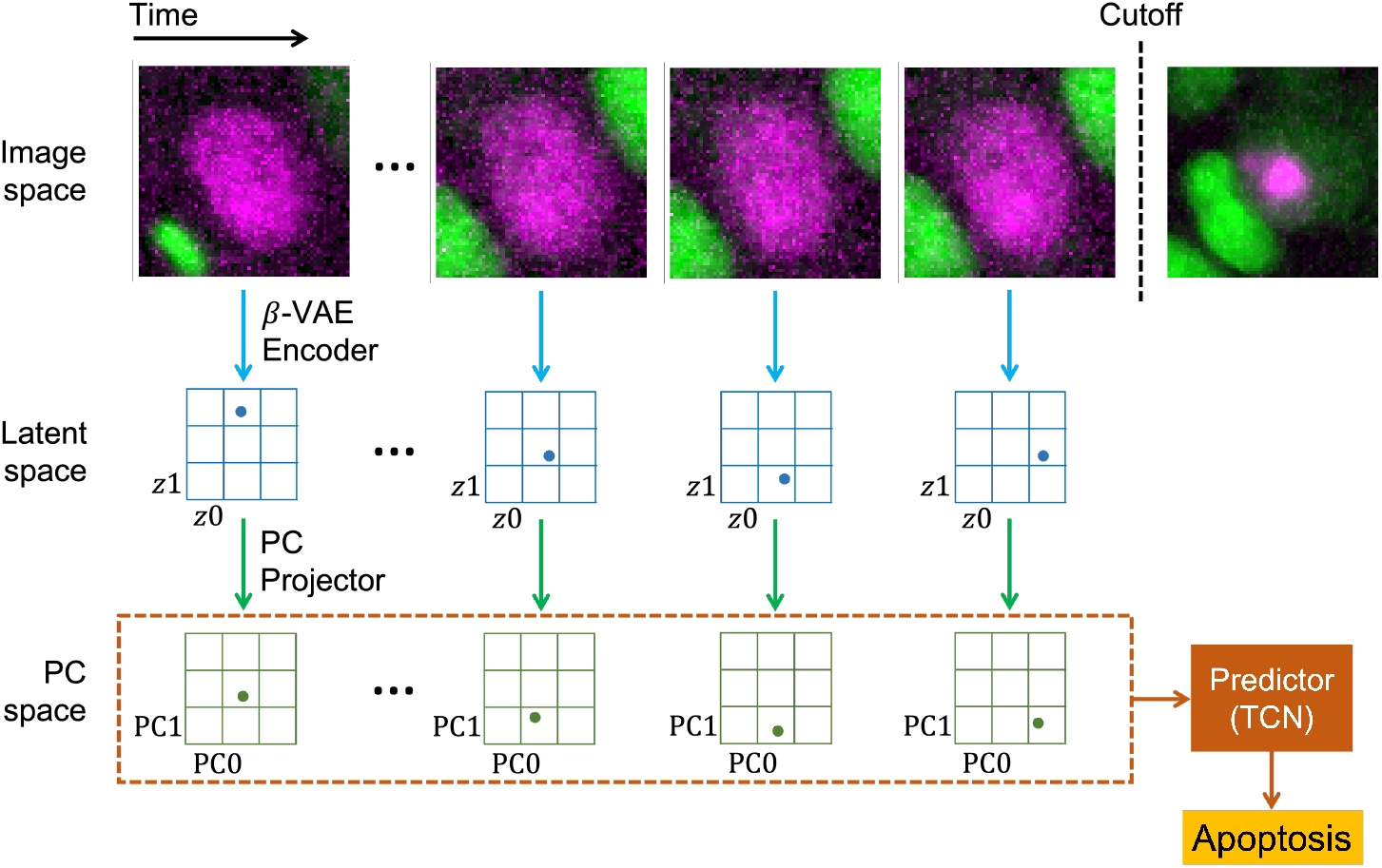
*τ* -VAE pipeline, from trajectory to prediction. The input is the trajectory in image space, trimmed to a cutoff such that it does not contain the fate event. Images are transformed into latent space by the *β*-VAE encoder, then into PC space by the PC projector, where they are collectively input to the TCN predictor.

The TCN was trained to predict scrib^kd^ cell trajectories as belonging to one of two potential fates: mitosis and apoptosis. A third “synthetic” class was added to represent those inputs that fall outside of the distribution (OOD) of real cell trajectories. Data for this class was artificially generated using a random-walk procedure in latent space [28]. The “synthetic” class was added to ensure that the *τ* -VAE learnt the predictors of both mitosis and apoptosis, rather than learning the predictors of one and “dumping” all remaining trajectories in the other fate class.

The PCs most important to cell fate prediction were then identified using an ablation procedure. This confirmed that the *τ* -VAE had learnt that nuclear area/local cellular density was the most important determinant of cell fate, completely independently of all the scientific studies that had arrived at the same conclusion.

The aim of the present study is to extend this work by characterising the impact of certain input features on the predictive behaviour of the *τ* -VAE. In particular, we sought to characterise the sensitivity of the model’s predictions to perturbations in both its internal representation of the input data, and perturbations in the input data itself.

## 3 Model perturbations

Perturbation is commonly used to uncover relationships between cause and effect. By varying one independent variable while keeping others constant, the specific contribution of that variable can be measured in a way that minimises exposure to confounding factors. This minimisation can be achieved by recording the statistically significant effects of a perturbation throughout a highly diverse population of inputs.

This logic allows us to examine the predictive behaviour of a deep neural network (or any other input-output model, in fact). By applying specific perturbations to the data representation at some point in the information processing pipeline, we can uncover causal relationships between model inputs and outputs. This can be either a physical perturbation, such as treating the cell culture with some drug, or a *virtual* perturbation, which involves altering the model’s internal representations of the underlying system without affecting the system itself (**Fig. 2**).

**Figure 2.**
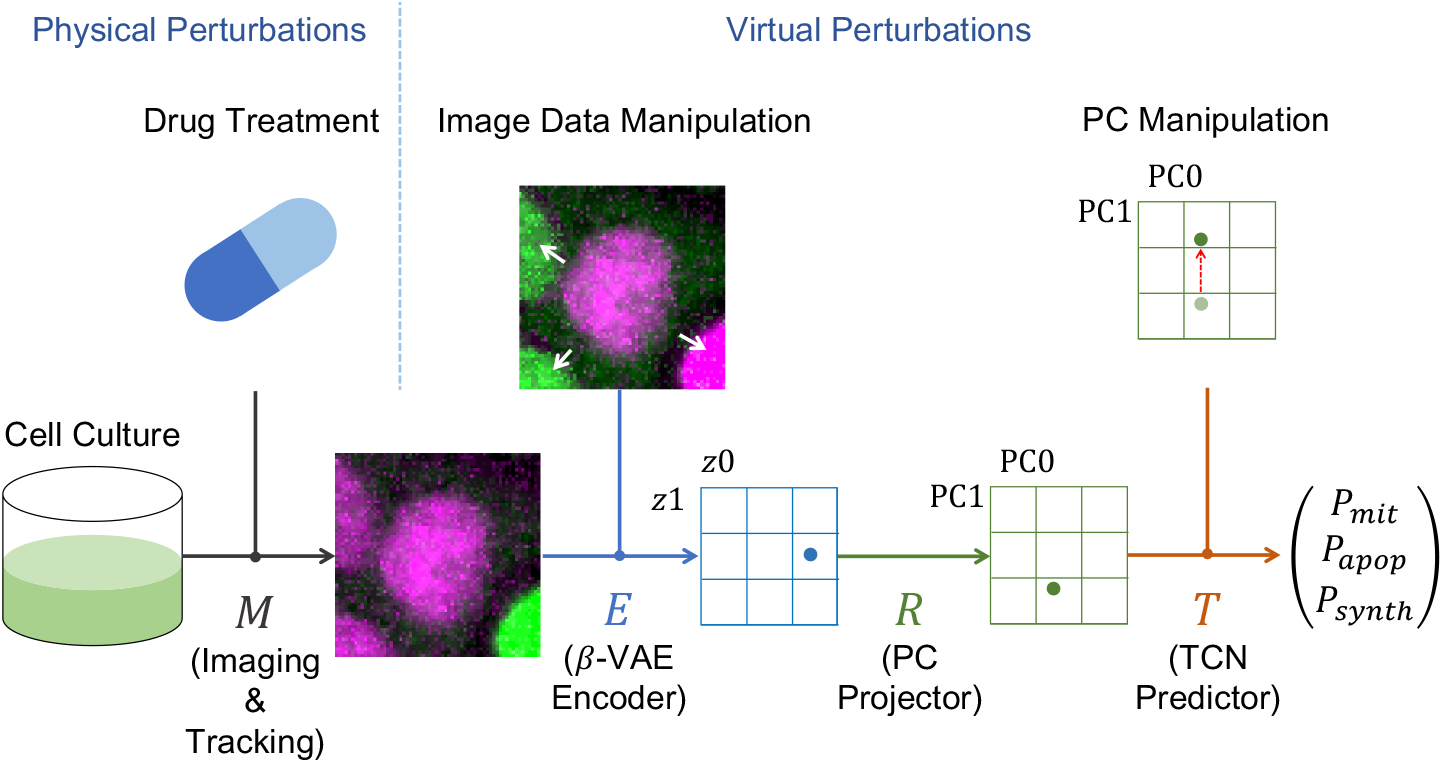
Virtual perturbation schemes. Location of the perturbations within the data pipeline. Neighbour cell type switching occurs before the encoding step (white arrows show perturbed cells). Nuclear area/PC1 adjustment occurs after PC projection. Drug treatments are applied directly to the cells before imaging (this perturbation does not feature in this work, but was used in our previous study [28]).

When applying a perturbation to the internal representation, it is often important that this representation is disentangled to a degree that enables the identification of particular latent features with specific physical/conceptual attributes, such as cell type or cell size. While achieving perfect disentanglement is extremely difficult, our *β*-VAE managed this to a high degree [28].

More generally, we can conceptualise a perturbation as some intervention applied to the data representation at some location in the pipeline (**Fig. 2**). In our case, the pipeline can be expressed as:

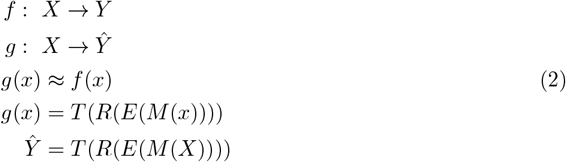

where *X* is the system under study (i.e., the cell competition), *M* is the imaging and tracking procedure, *E* is the *β*-VAE encoder, *R* is the PC projection and *T* is the TCN predictor. *Y* is then the true cell fate and 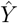is the predicted cell fate.

In this study, we focus on perturbations (denoted by Δ) applied to the time-lapse input data, post-tracking (Eq. 3), as well as on the model’s final internal representation of the input, in PC space (Eq. 4).

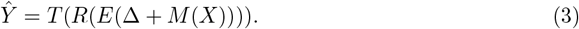

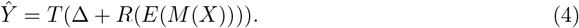

We applied perturbations that would allow us to further test the findings of our previous model and also represent situations that could plausibly occur in the underlying system, and so lie within the distribution of input trajectories used to train the *τ* -VAE (hereafter called the “realistic range”). So, for example, switching the cell type of neighbouring cells is a valid operation because it produces a situation that could plausibly occur in this range. Meanwhile, switching the cell type of the central cell is not valid because the *τ* -VAE was trained on scrib^kd^ cells and not MDCK^WT^ cells, hence switching the central cell type to MDCK^WT^ would push the perturbed trajectories outside the realistic range.

For the manipulation of input data, we chose neighbour cell type switching because it allowed us to test the conclusion of our own and other studies that neighbour cell type is not an important determinant of cell fate in scrib^kd^ cells [37, 38, 28]. Similarly, for the manipulation of internal representations, we decided to adjust PC1 because it allowed us to the test the finding that crowding/nuclear compression *is* an important determinant.

### 3.1 Manipulation of the input data

Wagstaff et al. [37] demonstrated that the presence of MDCK^WT^ cells induces apoptosis in scrib^kd^ cells only indirectly, by promoting greater crowding and a higher local cell density, which in turn mechanically compresses the scrib^kd^ cells. As the scrib^kd^ loser cells are less tolerant to crowding, they commit apoptosis. This suggests that the actual identity of the neighbours is not important, rather it is the degree of crowding/compression alone that determines the fate. Therefore, switching the cell type of neighbouring cells should have a small effect on cell fate prediction by the *β*-VAE, given that the perturbed trajectory should show similar degrees of compression to the original trajectory. The method we used to implement this was applied directly on the raw images, before they are input to the *β*-VAE encoder (**Fig. 2**). A binary map of cell locations was acquired using a U-Net based segmentation model [17]. This map was then used to alter the pixel values in each fluorescence channel such that GFP-tagged nuclei appear as RFP-tagged nuclei and vice versa (see Section 7.3 for details). After this step, the images are input through the three-stage *τ* -VAE as usual (**Fig. 3**).

**Figure 3.**
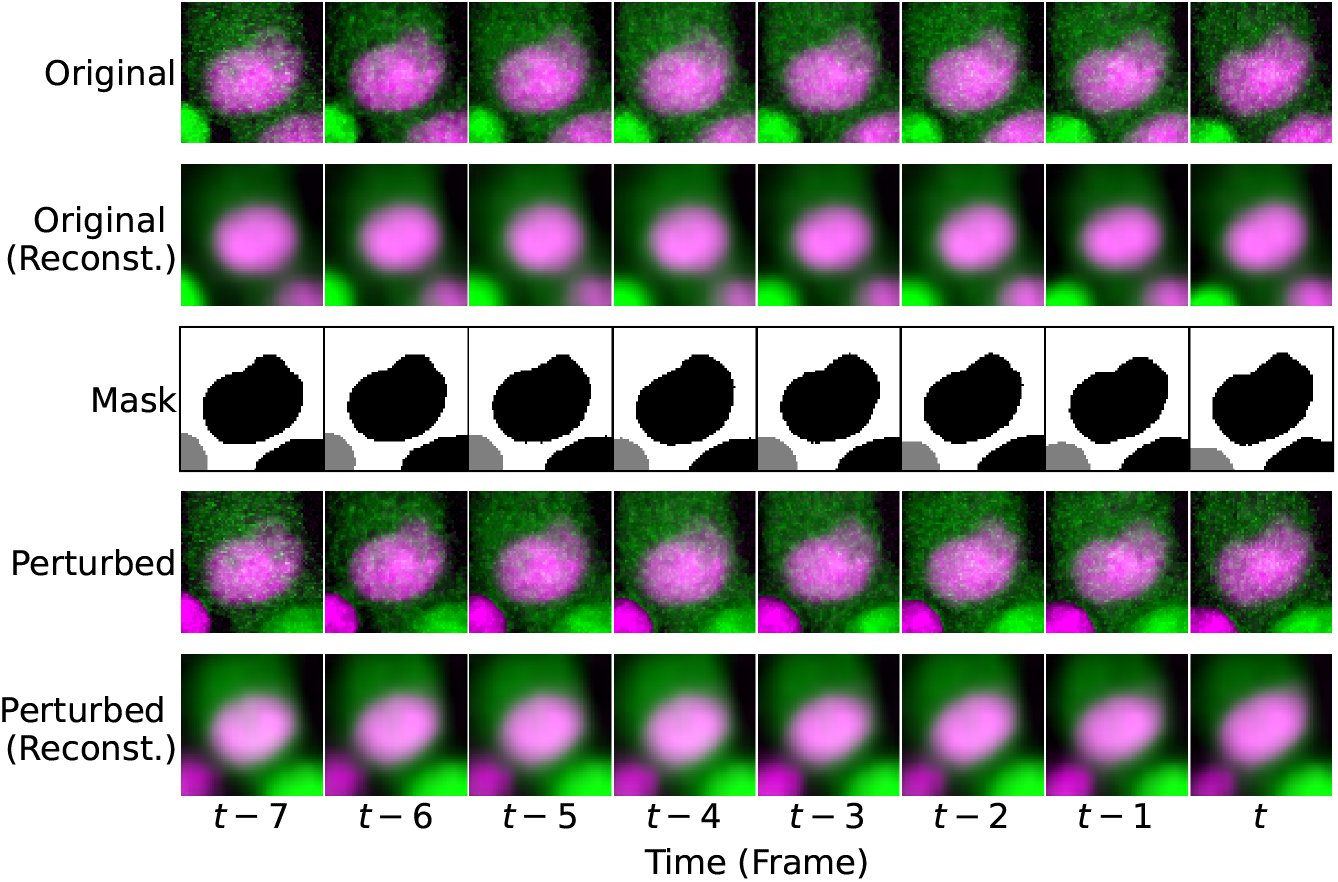
Neighbour cell-type switching procedure. Consecutive frames of one trajectory. In top-down order are shown: a) the original raw images, b) the *β*-VAE reconstructions of the original images, c) the U-Net derived segmentation masks (grey = MDCK^WT^, black = scrib^kd^), d) the perturbed images, and e) the *β*-VAE reconstructions of the perturbed images.

The predictive results of the *τ* -VAE on the perturbed dataset are shown in **Fig. 4**. Results were obtained by testing ten models on ten different testing sets (with *∼*300 scrib^kd^ cell tra-jectories each), obtained by 10-fold cross validation. Mean and standard deviation across the ten models were used to produce the confusion matrices (**Fig. 5**). The variation, compared to the unperturbed dataset, was very slight. There were small differences; in particular, the rate of mitosis prediction seemed to have increased marginally. This could indicate that the *τ* -VAE uses information about neighbour cell type to a small degree. However, it could also reflect the fact that when the cell type of neighbour cells is switched using our method, the *τ* -VAE’s representations of the *central* cell are also affected (as shown in **Fig. 3s**). This could in turn reflect the fact that certain configurations of cells are found only sparsely in the image dataset on which the *β*-VAE was trained.

**Figure 4.**
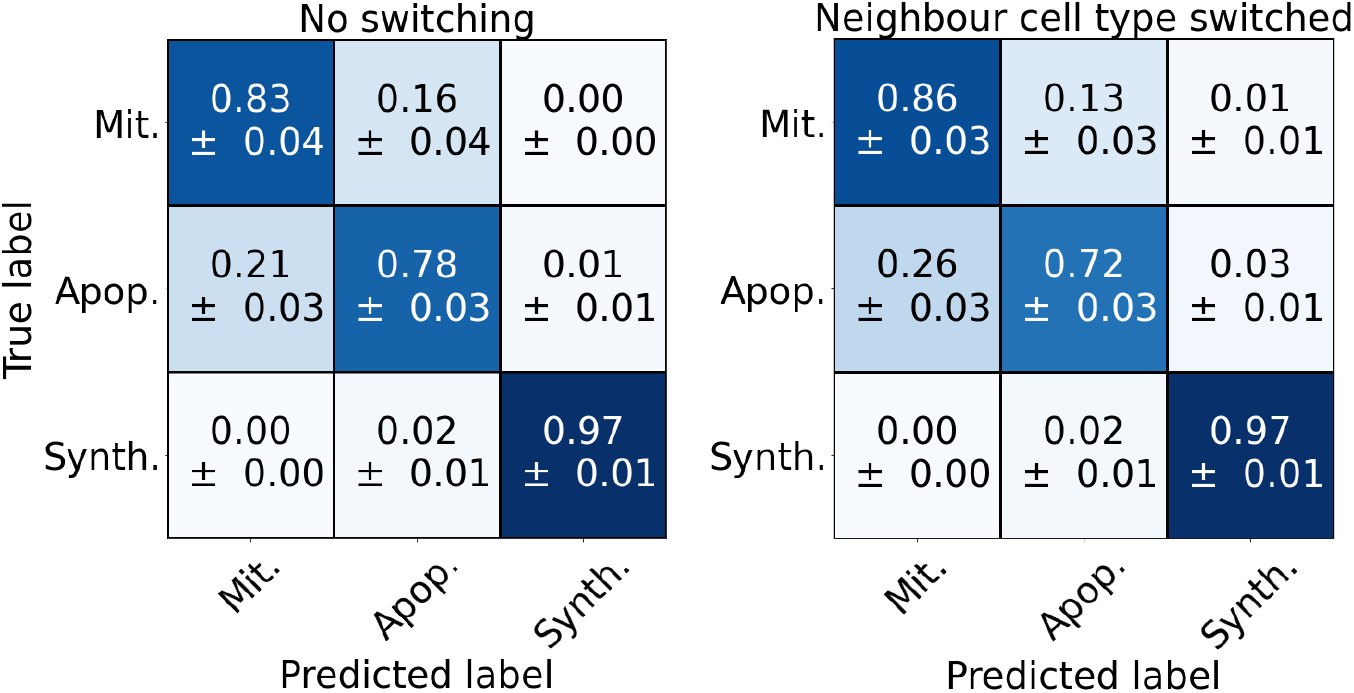
Confusion matrix showing the performance of the *τ* -VAE models on testing sets which have been perturbed by neighbour cell type switching. left) Confusion matrix on the un-perturbed dataset. right) Confusion matrix on the perturbed dataset, demonstrating that the identity of neighboring cells has little effect on the predicted cell fate.

**Figure 5.**
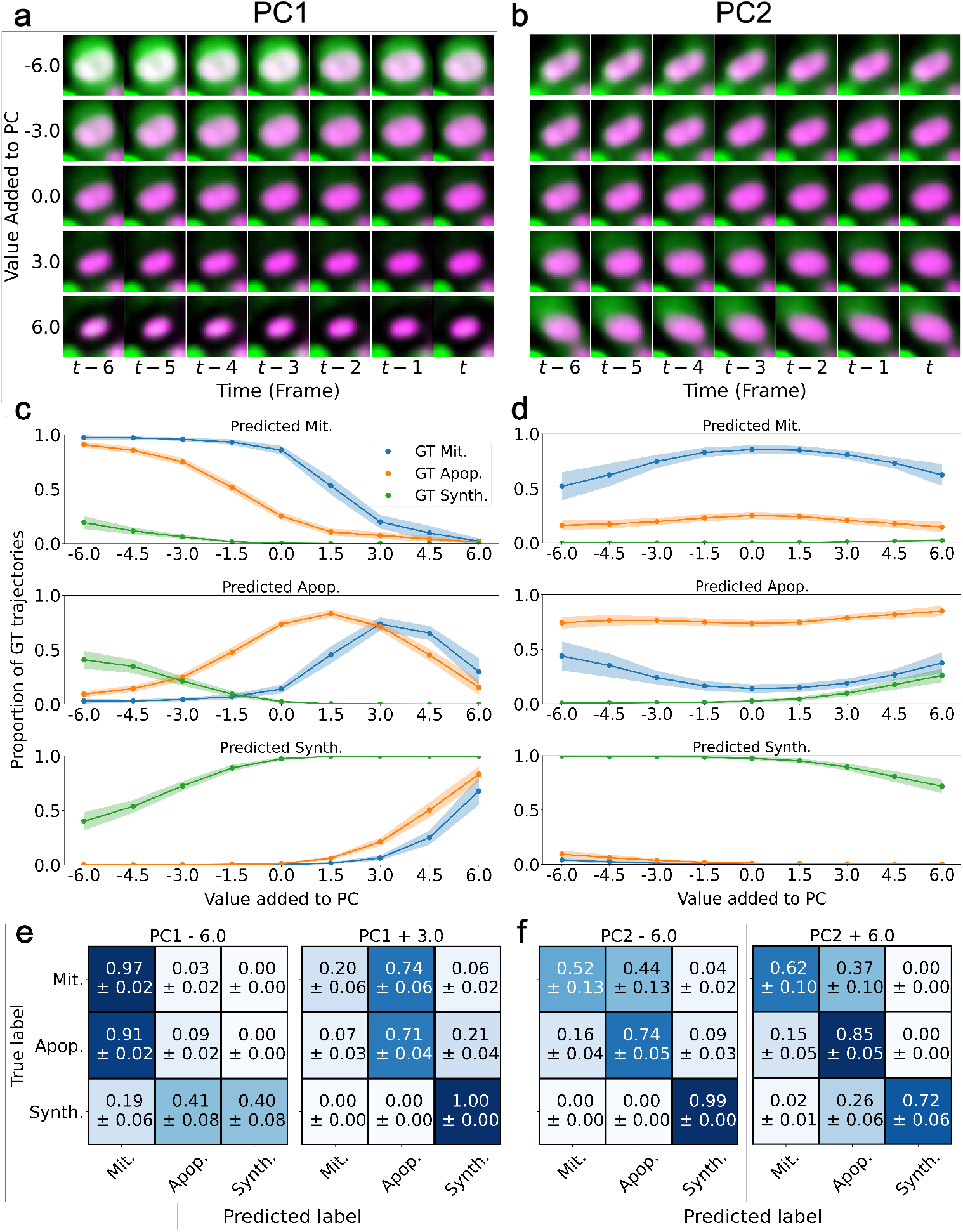
PC shifting. a, b) Decoded images, obtained by projecting the shifted PC representations back into latent space then passing the result through the decoder part of the *β*-VAE. Each row represents one constant value of a PC that has been added to each frame throughout the trajectory. c, d) Prediction outputs of the *τ* -VAE on the PC-shifted input data. Translucent fill-in indicates the standard deviation across models. e, f) Confusion matrices at both extremes of the realistic range for each PC.

Nevertheless, this perturbation step has demonstrated that the effect on predictor output of switching the cell type of neighbour cells is marginal at best, especially when compared to the large effects of adjusting PC1 - a perturbation whose consequences are of a similar magnitude in image space (Section 3.2).

As a further control, we tested the effects of switching the identity of the central cell (from scrib^kd^ to MDCK^WT^) rather than the neighbour cells. This led to the *τ* -VAE assigning a “synthetic” prediction to virtually all ground-truth apoptotic and mitotic trajectories. This result was expected due to the fact that the *τ* -VAE model under inspection was trained on real trajectories centred on scrib^kd^ cells only. Hence, MDCK^WT^ trajectories are out-of-distribution (OOD) with respect to the data on which the *τ* -VAE was trained, and the *τ* -VAE is evidently sensitive to this. OOD effects have implications for the scientific use of deep learning, as will be discussed later.

### 3.2 Manipulation of the internal representation

Mechanical compression of the nucleus, visually represented by a reduction of the nuclear area, has been shown experimentally to induce apoptosis in scrib^kd^ cells, even in the absence of MDCK^WT^ cells [37]. We therefore hypothesised that by virtually decreasing the area of the target cell nuclei in our images, we could increase the false-positive rate of apoptosis detection by our *τ* -VAE in a predictable manner.

The method we used to adjust nuclear area was to simply increase or decrease PC1, by a constant absolute value, throughout a trajectory (**Fig. 5a,c**). Since the TCN receives the PC representation of the trajectory, this step could be implemented prior to input to the TCN. Results were obtained by testing ten models on ten different testing sets, in the manner described in Section 3.1. Interestingly, the effect of adjusting PC1 is to simultaneously vary the nuclear area of the central cell, and the degree of crowding in the neighbourhood. This reflects the principle, learnt by the *β*-VAE, that higher local density of cells entails a higher degree of nuclear compression.

Virtual perturbations to PC1 induce a predictable, stable effect on the *τ* -VAE’s predictions. Increasing PC1 decreases the rate of mitosis prediction, while decreasing PC1 does the reverse. With increasing PC1, the rate of apoptosis prediction first increases, then decreases as the rate of synthetic prediction becomes dominant. This latter result shows that trajectories with such artificially high values of PC1 in fact do not lie within the distribution of real trajectories; in other words, within the “realistic range”. So, for trajectories which *do* lie within the distribution, the picture is one of monotonic decrease in mitotic probability and monotonic increase in apoptotic probability as PC1 is increased.

We have demonstrated that the specific adjustment of PC1 produces predictable and significant effects on the outputs of the *τ* -VAE. Our hypothesis is that these effects are due to the *nature* of the perturbation (nuclear area adjustment) and not simply its magnitude. To test this, we repeated the same procedure for the other thirty-one PCs aside from PC1.

In general, we did not see the same stable, predictable and monotonic relationship (within the realistic range) as with PC1. A case in point is PC2, whose adjustment induces a small rotation of the major axis of the central cell (**Fig. 5b**). Adjustment of PC2 produces very minor variations in the treatment of ground-truth apoptoses, and it triggers small decreases in performance in both the positive and negative direction for ground-truth mitoses (**Fig. 5d**). Evidently, the adjustment of PC2 is not sufficient to convert mitotic predictions to apoptotic predictions or *vice versa*.

The corresponding confusion matrices are shown in Fig. 5e. To demonstrate the effects of perturbation, we show predictions results for PC1 shifts of different sign. For PC1, we show the results of a +3.0 shift rather than a +6.0 shift because shifts greater than +3.0 tend to the push the trajectories outside of the realistic range, indicated by a large number of synthetic predictions. This is to ensure that we investigate only those trajectories that remain in the distribution of real-world data.

To quantify the relationship between *τ* -VAE output and each PC, we calculated the gradients of linear models fitted to the prediction curves by ordinary least-squares regression. The intuition is that PCs that monotonically changed the output prediction will show large gradients. In order to capture the PC-output relationship solely within the realistic range, we considered only those data-points for which less than 20% of ground-truth trajectories in a particular class were predicted as synthetic (indicative of the realistic range). As can be seen in Fig. 6, PC1 was associated with the greatest slope magnitudes, demonstrating that its relationship with model output was the most significant and monotonic. This suggests that certain physical features are important for cell fate determination (e.g. PC1/crowding) while some are not (e.g. PC2/rotation).

**Figure 6.**
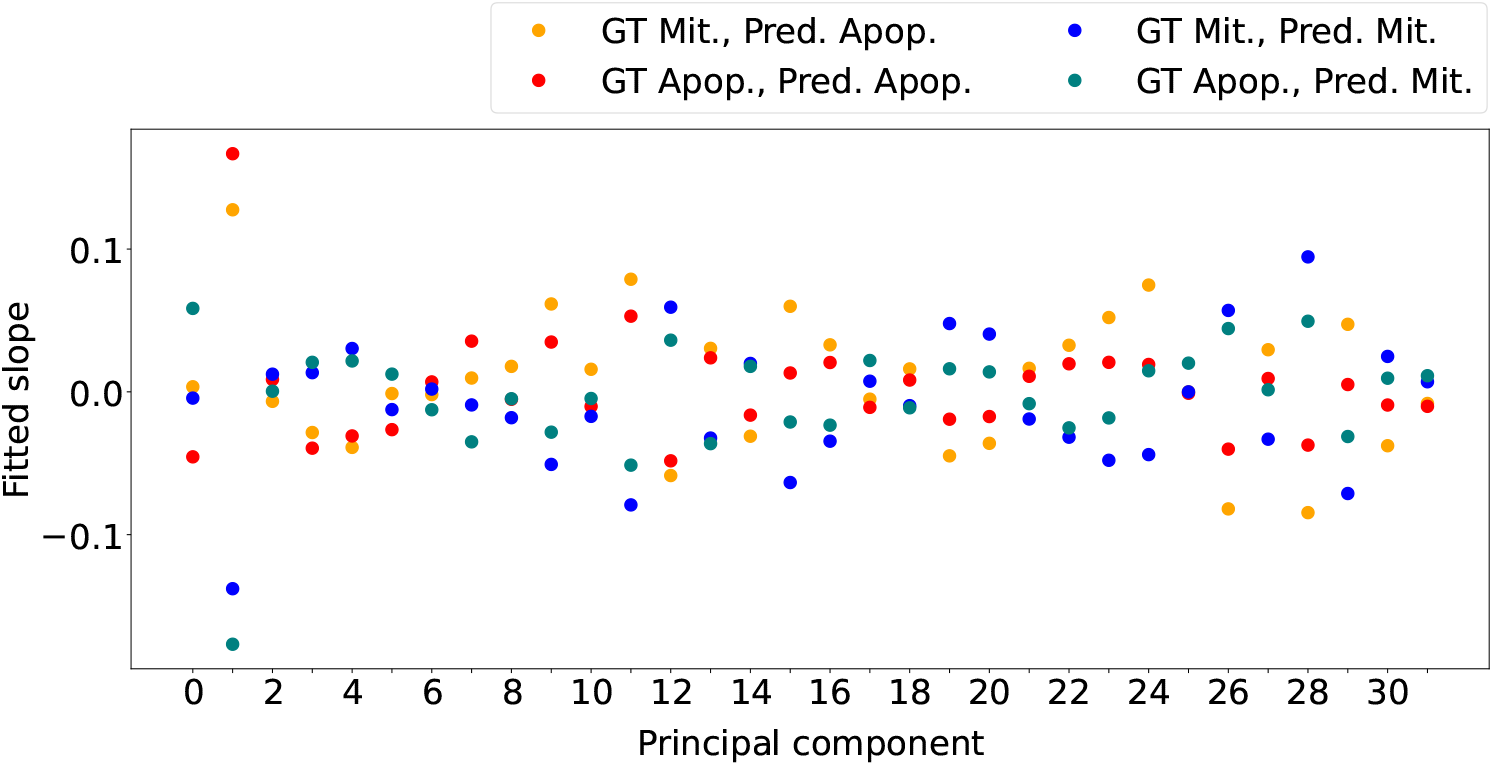
Slopes of linear models fitted to the prediction curves by ordinary least-squares regression. For each data-point, the mean value was considered. This calculation only used those data-points for which less than 20% of trajectories in the ground-truth class were predicted as synthetic, representing the “realistic range”.

## 4 Discussion

Virtual perturbation is a straightforward technique for assessing the impact of various input features on the output of a model, black box or otherwise. In this study we applied this method to a DNN trained to predict cell fate from image sequences, acquired in an experimental system that captures the phenomenon of mechanical cell competition. We show that altering nuclear area in the internal representation of the DNN exerts a significant, predictable and monotonic effect on the rate of apoptosis prediction within the realistic range. In contrast, flipping the cell type of neighbouring cells triggers only a minimal effect. Through this, we can confirm that our *τ* -VAE model has autonomously learnt that mechanical compression is a driving factor in apoptosis, while cell-type-dependent interactions with neighbours are not. Thus our deep-learning based approach has recovered the conclusions of previous experimental and statistical research on the system under study [37, 38].

Here, we explore only two types of virtual perturbation to our model system. One could imagine a multitude of others, each testing a different hypothesis. For example, by adjusting PC1 for varying lengths of time, instead of throughout the entire trajectory, one could investigate whether apoptosis is dependent on the duration of crowding, as well as its severity. The possibilities are limited only by our ability to apply perturbations in a controlled manner. That being said, it is important to consider whether the perturbations move the data out-of-distribution (OOD). DNNs trained to perform well within a specific distribution will not generally perform well outside of it. Hence, one should question the scientific value of perturbations that lead to OOD effects.

Moving forward, this technique could serve as a useful addition to the arsenal of deep-learning based scientific methods, which already includes feature ablation [28], linear discriminant analysis [27], symbolic regression [45, 46] and more. One advantage of virtual perturbation is that it does not require any assumptions regarding the form of the model or the form of the solution, in the way that linear discriminant analysis prescribes a linear classifier, and symbolic regression searches for elegant mathematical expressions. In fact, one advantage of this technique is that it allows for investigation of a model’s behaviour without detailed knowledge of its internal mechanisms. This is achieved by recording statistically significant consequences of perturbation across a population with as great a diversity as possible.

Moreover, we sought not only to test the resilience of a particular causal relationship throughout the dataset, but also across several different models, each trained to high accuracy. In Section 1.2 we introduced the notion of an equal-performance “Rashomon” set *G* [2]. The existence of multiple models in this set would indicate that there are multiple internal mechanisms through which high performance could be obtained, only one of which would correspond to the natural phenomenon *f*. We therefore sought to discover those input-output relationships that were robust across different members of *G*. We did this by training and testing ten different models through 10-fold cross-validation, reporting small standard deviations in our results across models (Figs. 4 & 5). Of course, it would be impertinent to say that this robustness automatically proves that the relationship extends to *f* as well; however, it certainly suggests that this is the case. In other words, our study certainly suggests that in the case of mechanical cell competition, crowding is a strong determinant of cell fate, on the basis that ten models independently trained to predict cell fate have all leveraged PC1 as an important input feature. Future studies could explore this concept in a more rigorous manner, to discover those properties common to all members of *G*.

Researchers in various domains may use virtual perturbation to generate hypotheses for physical experimentation. By rapidly exploring the space of possible experiments, it would allow researchers to carry out only those that have a good chance of being useful. For example, had there been no prior experiments on the effect of crowding on scrib^kd^ cell apoptosis, the present study would have flagged this as a hypothesis that could be tested. Overall, we hope that this technique will serve others well across a variety of domains, and that it will contribute meaningfully to the exciting field of deep-learning based scientific investigation.

## 5 Acknowledgements

We thank Nathan Day, Jasmine Michalowska and Dan Smaje for help annotating data, and Giulia Vallardi and Manasi Kelkar for additional supporting data. We thank Kristina Ulicna for feedback on the manuscript. We also thank members of the Lowe and Charras labs for discussions and technical support during the project.

## 6 Funding

This work was supported by a BBSRC LIDo AI PhD studentship to CJS. ARL wishes to acknowledge the Turing Fellowship from the Alan Turing Institute. ARL and GC wish to acknowledge the support of BBSRC grant BB/S009329/1.

## 7 Methods

For details on the data acquisition, image processing, as well as *β*-VAE and TCN training, and generation of synthetic trajectories, refer to Soelistyo et al. [28].

### 7.1 Principal component analysis

We used the PCA class from Scikit-learn (`sklearn.decomposition.PCA`) to extract 32 principal components (PCs) from our 32-dimensional latent space. The projective transformation that maps points in latent space to PC space consisted of the operation:

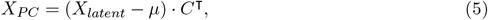

where *X*_*P C*_ and *X*_*latent*_ represent corresponding positional vectors in PC space and latent space, *μ* is a vector of per-component mean values, and *C* is a matrix of weights that define how each PC is calculated from each latent variable. The inverse projection is:

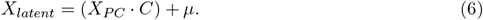

### 7.2 PC adjustment

Our method to adjust indiviudal PCs was to simply to add constant values to the PC of the encoding of each image across a trajectory. This step occurs after encoding by the *β*-VAE encoder and projection by the PCA model (Fig. 2). For a given trajectory with *T* time-points, and a given adjustment value of *v*, the transformation is:

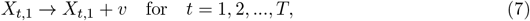

where *X* is a matrix of PC-space positional vectors; each row represents one time-point and each column represents one PC. To decode the PC shifted representations we first transformed the PC space vectors back into latent space using Eq. 6, then passed the result through the *β*-VAE decoder.

### 7.3 Cell type switching

To switch the cell type of neighbours, the U-Net segmentation mask was first used to determine the locations of the cells in an image. Then, in these locations, the maximum pixel value across both channels (GFP and RFP) was placed in the channel corresponding to the cell type opposite to the original cell type. We spared the central cell region from this operation by using the Scikit-image `morphology. label` function to distinguish the regions of each cell, then removing from the mask the region corresponding to the cell at the centre of the image. Given this final cell mask, the whole operation can then be described by:

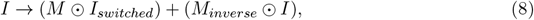

where *I* is the image, *M* is the binary mask denoting cell pixels, *I*_*switched*_ is a channel-switched representation of *I*, and *M*_*inverse*_ is the logical negation of *M*. The *⊙* operator is the Hadamard (element-wise) product.

Due to inherent intensity differences between the GFP and RFP signals, these were normalised throughout the original image on a per-channel basis prior to switching. We normalised each channel by first applying a median filter (`skimage.filters.median`) then removing outliers by detecting those pixels whose values deviated from the local median by some threshold. Then, we calculated the 5^*th*^ and 99^*th*^ percentile pixel values, divided the image by the absolute difference between these, and subtracted the 5^*th*^ percentile value from the result. Finally, we clipped the image to values between 0.0 and 1.0. The final, perturbed image was re-normalised such that each channel had a pixel value mean of 0.0 and standard deviation of 1.0 (this final normalisation step was done on all inputs to the *β*-VAE encoder, during both training and inference).

## Notes

### Competing Interest Statement

The authors have declared no competing interest.

